# Deficits in tail-lift and air-righting reflexes in rats after ototoxicity associate with loss of vestibular type I hair cells

**DOI:** 10.64898/2026.03.24.712950

**Authors:** Aïda Palou, Michele Tagliabue, Mathieu Beraneck, Jordi Llorens

## Abstract

The rat vestibular system plays a critical role in anti-gravity responses such as the tail-lift reflex and the air-righting reflex. In a previous study in male rats, we obtained evidence that these two reflexes depend on the function of non-identical populations of vestibular sensory hair cells (HC). Here, we caused graded lesions in the vestibular system of female rats by exposing the animals to several different doses of an ototoxic chemical, 3,3’-iminodipropionitrile (IDPN). After exposure, we assessed the anti-gravity responses of the rats and then assessed the loss of type I HC (HCI) and type II HC (HCII) in the central and peripheral regions of the crista, utricle and saccule. As expected, we recorded a dose-dependent loss of vestibular function and loss of HCs. The relationship between hair cell loss and functional loss was examined using non-linear models fitted by orthogonal distance regression. The results indicated that both the tail-lift reflex and the air-righting reflexes mostly depend on HCI function. However, a different dependency was found on the epithelium triggering the reflex: while the tail-lift response is sensitive to loss of crista and/or utricle HCIs, the air-righting response rather depends on utricular and/or saccular integrity.

## 1. INTRODUCTION

The vestibular system in the inner ear is a transducer of angular and linear accelerations, including gravity (reviewed by Cullen, 2023; Curthoys et al., 2017; Eatock and Songer, 2011). On each side of the mammalian head, five sensory epithelia contain the transducing cells, named hair cells (HC). There are two types of HC, type I (HCI) and type II (HCII). The five epithelia are three cristae at the end of their corresponding semi-circular canals, primarily encoding angular accelerations, and two macula, utricle and saccule, mainly encoding linear accelerations. In each of these receptors, several histological, molecular and functional criteria allow to define a central zone (also called striola zone in the utricle and saccule) and a peripheral zone. Vestibular information serves to drive the vestibulo-ocular and vestibulo-spinal stabilizing reflexes, and has also important roles in cognitive function such as spatial orientation and navigation.

Vestibulo-spinal reflexes are well-known contributors to equilibrium and motor performance (Cullen, 2023). Accordingly, lesions of the peripheral vestibular system result in overt motor deficits, such as the loss of the tail-lift and the air-righting reflexes in rodents. Thus, vestibular deficient rats and mice display a trunk flexion response instead of the normal extension reflex during a tail-lift manoeuvre (Hunt et al., 1987; Llorens et al., 1993; Soler-Martín et al., 2007; Martins-Lopes et al., 2019) and fail to right themselves in the air as healthy rodents do when dropped supine onto a foam cushion (Pellis et al., 1989; Shoham et al., 1989; Ossenkopp et al., 1990; Llorens et al., 1993; Soler-Martín et al., 2007; Martins-Lopes et al., 2019). At present, the knowledge on the effects of partial vestibular lesions on these reflexes is incomplete and further studies are needed for at least two reasons. First, partial vestibular lesions are common in clinical settings, as a result of ototoxicity, ageing or other conditions. Therefore, a detailed knowledge on the relationship between the extent of the vestibular lesions and the extent of the alteration of the vestibular reflexes should ease the understanding of the physiopathological basis of clinically recorded deficits. Second, these relationships may also shed light on the differential role of the different parts of the vestibular system (i.e. HCI vs HCII, central vs. peripheral zones, crista vs macula) on each function. These interests are well illustrated by our studies on the impact of partial or reversible vestibular dysfunction on gaze control in mice (Schenberg et al., 2023, 2026). These studies revealed that HCI have a greater role than HCII in the vestibulo-ocular reflex and that transient vestibular loss causes a change in weight in the use of vestibulo-ocular and optokinetic reflexes for gaze stabilization that persists after vestibular repair, a conclusion with significant clinical value.

To study vestibulo-spinal reflexes in rats, we developed quantitative measures of the tail-lift reflex and the air-righting reflex by measuring the tail-lift angle and the air-righting time in high speed video recordings (Martins-Lopes et al., 2019; Maroto et al., 2021a,b). In one study (Maroto et al., 2021a), we examined the relationship between reflex loss and extent of lesion in the vestibular epithelium and reached the conclusion that these two reflexes depend on the function of non-identical populations of HCs. Now, we are presenting the results of a new assessment of this relationship, which was deserved for several reasons. First, the previous study (Maroto et al., 2021a) included some uncertainties on the identification of HCI and HCII. Now, the identification of new molecular markers, particularly osteopontin (known also as SPP1) as HCI marker (McInturff et al., 2018; Borrajo et al., 2024), and the availability of a detailed description of the HC type composition of the vestibular epithelia (Borrajo et al., 2024) allow for a more robust analysis of the pathology. Second, this previous study included only male rats. This choice was driven by the fact that male and female rats show a measurable difference in the dose-effect relationship of the vestibular toxicity of the compound used for causing the lesions, 3,3’-iminodipropionitrile (IDPN). For this reason, only rats of the better described sex (males) were used in the previous study and the characterization of females was pending. Therefore, in the present study we used female rats and included osteopontin as a positive HCI marker. Finally, the present study includes a different mathematical analysis of the data that provides a deeper insight on the relationship between the sensory epithelium damage and the vestibular function loss.

## 2. METHODS

### 2.1. Animals and treatments

The use of animals in this research was done in accordance with EU Directive 2010/63, as implemented by Law 5/1995 and Act 214/1997 of the Generalitat de Catalunya, and Law 6/2013 and Act 53/2013 of the Gobierno de España, and approved by the Ethics Committee on Animal Experimentation of the Universitat de Barcelona (protocol number 155/21 (P2)). A total of 38 young adult female Long-Evans rats (175-199 g) were purchased from Janvier Labs (Le-Genest-Saint-Isle, France) and housed for six days of acclimatization before the experiment began. Housing conditions were: two animals per cage; standard cages (215 ×465 ×145 mm) with wood shavings as bedding; wood sticks and cardboard shavings as enrichment elements; 12:12 h light:dark cycle (lights on at 7:30 am); temperature of 22 ± 2 °C; food (TEKLAD 2014, Harlan Laboratories, Sant Feliu de Codines, Spain) and tap water ad libitum. Rats were regularly weighed and evaluated for overall toxicity to limit suffering according to ethical criteria during the experiments. No animals had to be euthanised or died after the treatment.

Rats were distributed into 6 treatment groups, receiving 0 (vehicle control group, n=6), 150 (IDPN_3X150, n=6), 175 (IDPN_3X175, n=6), 200 (IDPN_3X200, n=6), 225 (IDPN_3X225, n=6), 250 (IDPN_3X250, n=4) or 300 (IDPN_3X300, n=4) mg/kg per day for 3 consecutive days of IDPN (CAS # 111-94-4, > 98%, cat. # I0010 - TCI Europe, Zwijndrecht, Belgium). IDPN was administered i.p. in 2 ml/kg of saline. This compound causes vestibular HC degeneration in rats and other mammalian and non-mammalian vertebrates (Llorens et al., 1993; Soler-Martin et al., 2007; Wilkerson et al., 2018).

Animals were euthanized at days 26-28 after exposure. Their temporal bones were quickly collected on cold 4% paraformaldehyde in phosphate buffered saline (PBS) and the vestibular epithelia obtained using a binocular stereomicroscope under a fume hood. After collection, epithelia were fixed for 1 h in 4% paraformaldehyde in PBS, and then stored at −20°C in a cryoprotective solution (34.5% glycerol, 30% ethylene glycol, 20% PBS, 15.5% distilled water) until further processing.

### 2.2. Assessment of vestibular reflexes

The tail-lift reflex and the air-righting reflex were evaluated before IDPN exposure (pre-test, Day 0) and at days 1, 4, 7, 14, 21 and 25 after the day of the last IDPN dose. The reflexes were assessed as described in detail elsewhere (Martins-Lopes et al., 2019; Maroto et al., 2021a,c). Briefly, for the tail-lift reflex, the rat is gently pulled up by the base of the tail until lifted to approximately 40 cm and then immediately returned down using a smooth and continuous movement. The response of the animal to the tail-lift manoeuvre is video recorded from the side at high speed (240 frames/s). Healthy rats show an anti-gravity or landing reflex of limb and trunk extension, while vestibular deficiency results in ventral flexion of the trunk. The movies are then used to assess the minimum angle formed by the nose, the back of the neck and the base of the tail during the manoeuvre. For the air-righting reflex, the experimenter holds the rat supine at approximately 40 cm above a foam cushion and suddenly releases it. Healthy rats quickly right themselves in the air, while vestibular deficiency results in delayed or absent righting. The reflex is recorded also at 240 fps from the front and the movies used to assess the time to right of the rat’s nose. We used a GoPro Hero 5 camera for recording and the Kinovea software (www.kinovea.org) for video analysis as detailed in Martins-Lopes et al. (2019).

### 2.3. Immunohistochemistry

Vestibular epithelia were immunolabelled following the protocol of Lysakowski et al. (2011) as described in detail elsewhere (Maroto et al., 2021a, b). Briefly, samples were rinsed in PBS, permeabilized and blocked with 4% Triton-X-100 and 20% donkey serum in PBS for 1h, incubated with the primary antibodies in PBS with 0.1% Triton-X-100 and 1% donkey serum for 24h, incubated with the secondary antibodies in PBS with 0.1% Triton-X-100, and whole-mounted in Fluoromount (Sigma-Aldrich cat# F4680). The mixture of primary antibodies included (1) mouse monoclonal anti-MYO7A (Developmental Studies Hybridoma Bank cat# 138-1-s, RRID: AB_2282417; 1/100); (2) guinea pig anti-calretinin (Synaptic Systems cat# 214.104, RRID: AB_10635160; 1/500); (3) goat anti-osteopontin/SPP1 (RD Systems cat# AF808, RRID: AB_2194992; 1/400); and (4) rabbit anti-oncomodulin (Swant cat# OMG4, RRID AB_10000346; 1/400). Secondary antibodies were donkey Alexa-647 anti-mouse IgGs (Invitrogen cat# A31571; 1/500), Alexa-555 anti-goat IgGs (Invitrogen cat# A21432; 1/500), Alexa-488 anti-guinea pig IgGs (Jackson ImmunoResearch cat# 706-545-148; 1/500), and Daylight-405 anti-rabbit IgGs (Jackson ImmunoResearch cat# 711-475-152; 1/200).

### 2.4. Epithelial imaging and cell counts

After fluorescent immunolabeling, the vestibular epithelia were imaged using a LSM880 Zeiss confocal microscope with a 63x objective (numeric aperture: 1.4). Oncomodulin label was used to identify the central/striola region of the receptors and Z-stacks of 0.5 μm were obtained from square areas (67.5 × 67.5 µm) of the central and peripheral zones of the three vestibular organs (horizontal semi-circular canal, utricle and saccule). In the utricle and saccule periphery, a medial / internal zone and a lateral / external zone were defined with respect to the striola as shown in the schemas included in the figures. The images were obtained from the approximately same location within the epithelium in all animals. The ImageJ software (National Institute of Mental Health, Bethesda, Malyland, USA) was used for cell counting. In each Z-stack image, the total number of HCs was obtained by counting the number of cells labelled with the anti-MYO7A antibody (Hasson et al., 1997; Borrajo et al., 2024). HCI were identified by anti-osteopontin/SPP1 label, particularly intense in the neck region of the cell (McInturff et al., 2018; Borrajo et al., 2024). In the striola/central region of the organs, we also determined the number of HCI encased by calyces from calyx-only afferents that express calretinin (Desmadryl and Dechesne, 1992; Maroto et al., 2021b). To estimate the number of HCII, we counted HCs expressing calretinin (Dechesne et al., 1991). Calretinin is expressed by 85-90% of the HCII and is not expressed by HCI, and the 10-15% of HCII that do not express calretinin represent a 4-5% of the total population of HCs (Borrajo et al., 2024).

### 2.5. Data analysis

Data analyses were performed using the IBM SPSS Statistics 25 or the GraphPad Prism 10 program packages. Body weight, tail lift angle and air-righting time data were analysed by repeated measures MANOVA with time as the within-subject factor. Significant treatment by day interaction was followed by day-by-day ANOVA analyses and Duncan’s post-hoc tests. Dose-dependency assessment of selected variables was evaluated by one-way ANOVA analysis and Tukey’s multi-comparison test. To characterize the relationship between HC counts and behavioural measures, data were normalized as follows: 1) The HC remaining in the epithelia was expressed as the percentage of mean count of the corresponding cell type and epithelial region in the control group; 2) Behavioural data were expressed as the percentage between the values defined by the mean values of the control and high-dose (IDPN_3x300) groups, set as 100% and 0%, respectively. After normalization, pairs of residual HC and behavioural data were fitted using two-dimensional non-linear (sigmoidal) models, with parameter estimation performed by orthogonal distance regression. The local slope of each fitted sigmoid was then computed across the full range of HC loss, and regions of “moderate” slope were defined as those where the slope angle fell between 10° and 80°. Slopes below 10° (approaching 0°) correspond to HC loss occurring without a noticeable change in reflex performance. Conversely, slopes above 80° (approaching 90°) correspond to a decline in reflex performance that is not accompanied by HC loss. A moderate slope (10° to 80° range) therefore reflects a functional association between HC loss and functional impairment. The extent of the HC loss range over which the sigmoid slope remained moderate was called Functional Cell Range (FCR) and was used as a quantitative indicator of how closely the changes in reflex performance tracked HC loss.

## 3. RESULTS

### 3.1. Effects of IDPN on vestibular function and vestibular hair cells

IDPN caused dose- and time-dependent effects on body weight and vestibular function. Transient decreases in body weight were observed in all treated groups but the effect in IDPN_3X150 rats was small (less than 2% decrease) and disappeared by day 2 after the last of the three dosing days. In the other threated groups, the maximal effect was recorded at day 4 post-dosing, with 8-10 % decrease in the three high dose groups. After this day, body weights returned to normal daily increases. The loss of vestibular function manifested in all groups of animals treated with IDPN_3X175 or higher doses as a syndrome of abnormalities in spontaneous motor behaviour and an impairment in anti-gravity reflexes (Fig. 1), including the landing reflex in response to a lift by the tail, and the righting reflex during a fall initiated in supine position, as described in previous studies in male rats (Llorens et al., 1993; Llorens and Rodríguez-Farré, 1997; Martins-Lopes et al., 2019; Maroto et al., 2021a,c). Rats exposed to the lowest dose of IDPN (3 × 150 mg/kg) showed control-like reflexes. In the affected animals, the minimum angle measured between the nose, the back of the neck and the base of the tail during the tail-lift manoeuvre decreased and attained a minimum value at day 7 after dosing, with higher doses inducing a faster decline in the value (Fig 1A). Animals treated with IDPN_3X225 or higher doses showed a maximal effect and no recovery. An almost maximal effect was recorded in the IDPN_3X200 animals on day 7 followed by an only partial recovery, and a complete recovery from an intermediate effect was observed in the IDPN_3X175 animals. Repeated-measures MANOVA of the angles resulted in significant day (F[6,26]=48.1, p<0.001), treatment (F[6, 31]=32.8, p<0.001) and day X treatment (F[36, 116.9]=5.39, p<0.001) effects. Day by day ANOVA analyses were significant from day 4 to 25 (all F’s [6, 31] > 14.7, all p’s < 0.001). IDPN also caused an increase in the time required to right in the air (Fig 1B), a recognized consequence of vestibular function loss (Ossenkopp et al., 1990). In this case, the dose-response relationship was shifted to the right in comparison to that of the tail-lift reflex. Thus, no significant effects were recorded in the IDPN_3X150 and IDPN_3X175 groups, and a transient effect with complete recovery was observed in the IDPN_3X200 group of rats. On day 25 after dosing, only the three groups with the highest doses of IDPN (3X225, 3X250 and 3X300) showed increased times in the air-righting reflex in comparison to mean time of the control group. Repeated-measures MANOVA of the times resulted in significant day (F[6,26]=64.6, p<0.001), treatment (F[6, 31]=39.6, p<0.001) and day X treatment (F[36, 116.9]=6.36, p<0.001) effects. Day by day ANOVA analyses were significant from day 4 to 25 (all F’s [6, 31] > 9.18, all p’s < 0.001). To measure the final effect of the IDPN exposure on the vestibular reflexes, we calculated the mean tail-lift angle and air-righting time values of the two last days (21 and 25) of assessment after dosing (Fig 1C, D). ANOVA analyses of these data resulted in significant effects of IDPN on tail-lift angles (Fig 1C - F[6,31]=35.11, p<0.0001) and on air-righting times (Fig 1D - F[6,31]=41.03, p<0.0001). Post-hoc multiple comparisons analyses revealed significant differences with respect to the control group in doses of IDPN of 3x 200 mg/kg or higher for the tail-lift test and of 3x 225 mg/kg or higher for the air-righting test.

**Figure 1.**
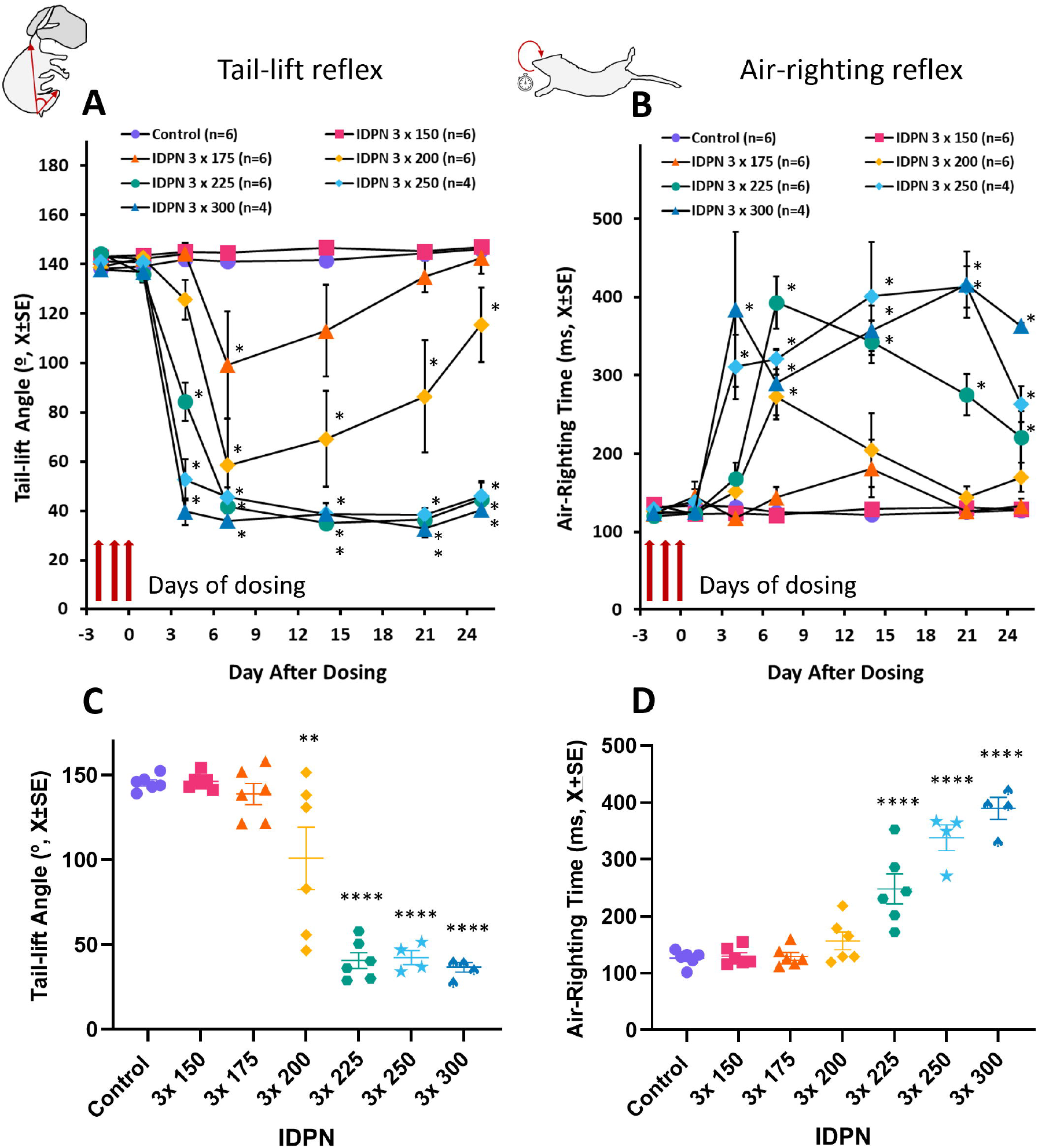
Effects of 3,3’-iminodipropionitrile (IDPN) on vestibular reflexes in adult female Long-Evans rats. Animals received one i.p. injection per day of IDPN (0 [Control] to 300 mg/kg/day) for 3 days. **A, C.** Tail-lift reflex: minimum nose-neck-base of the tail angles displayed by the rats when lifted by the tail and lowered back. **B, D**. Time course of the air-righting reflex: righting times displayed by the rats when dropped in supine position from approximately 40 cm above a foam cushion. **A, B**. Time course; data are mean + SE values. *: p<0.05, significantly different from control group, Duncan’s test after significant ANOVA and repeated-measures MANOVA analyses. **C, D**. Final effect; each data point is the average of angle (C) or time(D) values at days 21 and 25 after exposure of an individual animal. Lines are mean ± SE values per dosing group. **: p<0.01, ****: p<0.0001, significantly different from control group, Dunnett’s multiple comparisons test after significant ANOVA.

IDPN also caused a dose-, type- and region-dependent loss of HC in the vestibular epithelia, as assessed at 26-28 days after exposure by immunofluorescent labelling and confocal microscopy imaging (Fig 2, 3 and Supplementary Fig S1-S3). Figure 2A shows representative images of the overt effect of IDPN on HC density as a function of the dose. For a detailed evaluation of the effect, we counted, for each animal, receptor and zone, the number of total HCs (MYO7A+), number of HCIs (SPP1+), number of HCs in the main (calretinin+) HCII population, and number of calyx-only HCIs (SPP1+ surrounded by a calretinin+ calyx). The numbers of total HCs, shown in Figure 2B, reflects the region-dependency of the IDPN effect, as the dose-response relationship differs depending on the receptor and zone being considered. Thus, the lowest effective dose was that of the IDPN_3X175 animals, that caused a modest loss of HCs in the periphery of the crista and medial periphery of the utricle and a deeper loss of HCs in the centre of these receptors. In contrast, neither this dose nor the following one (3X200) caused a significant effect in the saccule, where the IDPN_3x225 group was the first one displaying significant loss of HCs.

**Figure 2.**
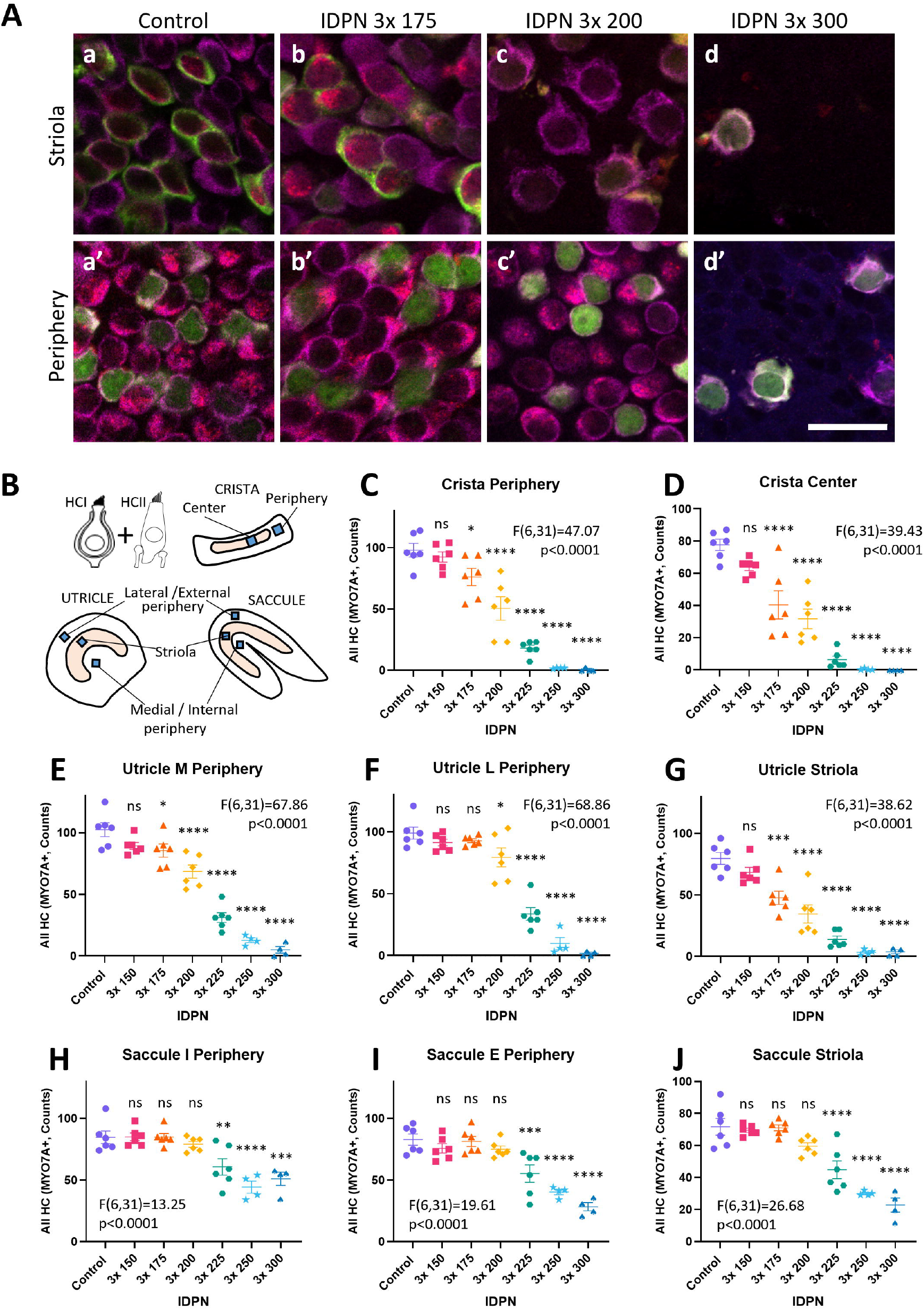
Effects of 3,3’-iminodipropionitrile (IDPN) on the vestibular sensory epithelium in adult female Long-Evans rats. Animals received one i.p. injection per day of IDPN (0 [Control] to 300 mg/kg/day) for 3 days. **A.** Example images of the effect of IDPN on the density of hair cells (HCs) in the sensory epithelium. Examples are from the striola region (upper row) and the lateral periphery (lower row) of the utricle. Overlay images are shown with MYO7A (magenta), osteopontin/SPP1 (red), and calretinin (green) labels. Scale bar = 20 µm. **B**. Schematic drawing of the two types of HC (HCI and HCII), the three organs (crista, utricle and saccule), and the regions sampled in the organs (center and periphery in the crista; striola, lateral periphery and medial periphery in utricle; striola, external periphery and internal periphery in saccule). The blue squares show the approximate region where the images were obtained for counting. **C-D**. Number of total HCs (HCI+HCII) obtained counting the number of MYO7A+ profiles per image in each organ and region. Points and lines display individual and mean ± SE values per dosing group, respectively. F and p values are from one-way ANOVA analyses. ns: non-significant, *: p<0.05, **: p<0.01, ***: p<0.001, ****: p<0.0001, compared to control group, Dunnett’s multiple comparisons test after significant ANOVA.

**Figure 3.**
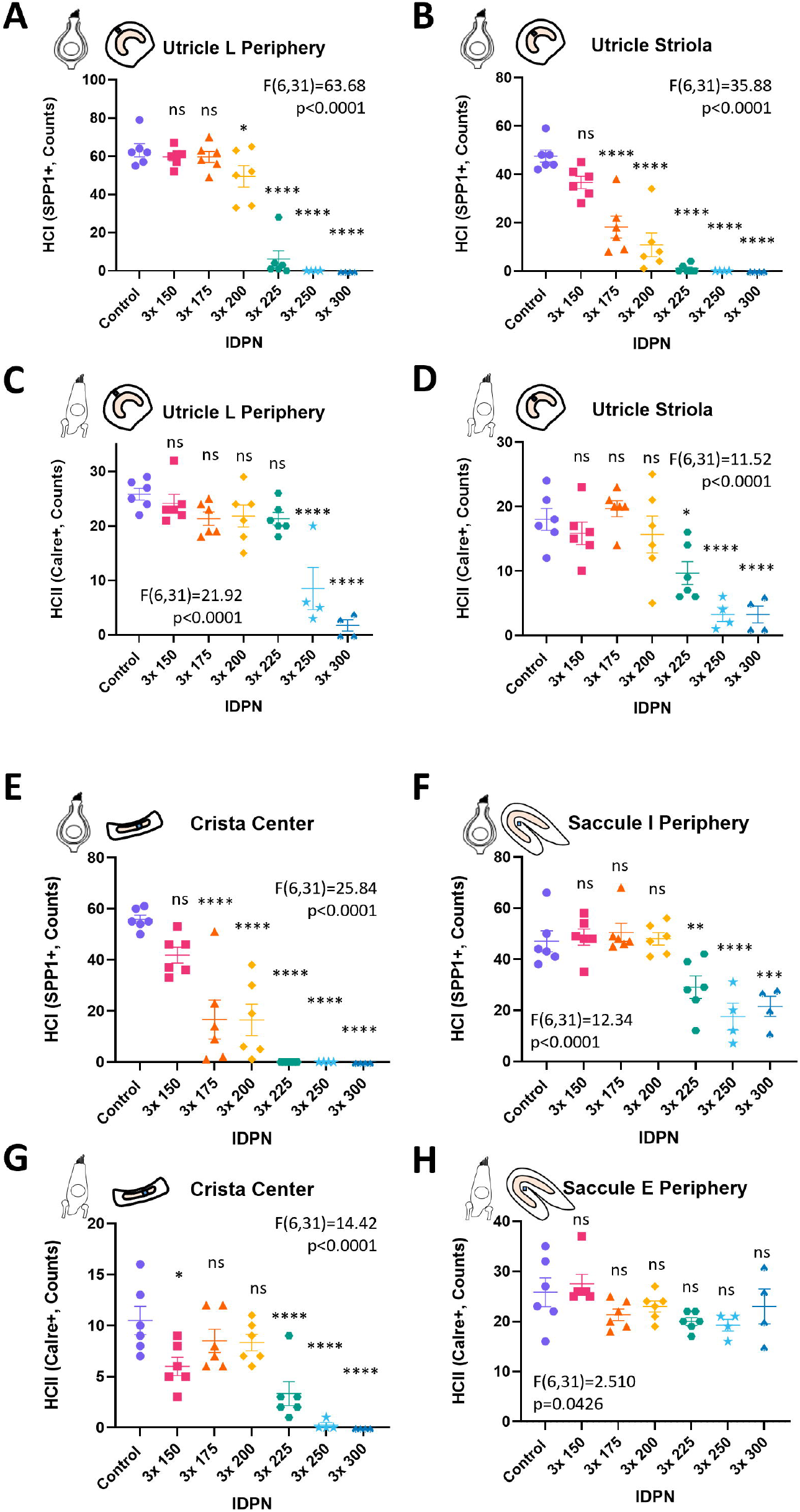
Representative examples of the dose-response effect of 3,3’-iminodipropionitrile (IDPN) on the counts of HCI and HCII in the vestibular organs of adult female Long-Evans rats. Animals received one i.p. injection per day of IDPN (0 [Control] to 300 mg/kg/day) for 3 days. **A-D.** Utricle data allowing to compare the dose-response relationships between HCI **(A, B)** and HCII **(C, D)** in the lateral periphery **(A, C)** and the striola **(B, D)** regions. E-H. Comparison of HCI **(E, F)** and HCII **(G, H)** dose-response data between an area highly sensitive to ototoxic damage (crista center: **E, G)** and an area of low sensitivity (saccule internal periphery: **F, H)**. HCI and HCII counts correspond to HC expressing osteopontin or calretinin label, respectively. Points and lines display individual and mean ± SE values per dosing group, respectively. F and p values are from one-way ANOVA analyses. ns: non-significant, *: p<0.05, **: p<0.01, ***: p<0.001, ****: p<0.0001, compared to control group, Dunnett’s multiple comparisons test after significant ANOVA.

As expected, the analysis of cell counts according to IDPN dose, cell type and epithelial location indicated that different cell populations showed different sensitivity to the IDPN-induced loss. Supplementary Figures S1 – S3 show complete dose-response data for all epithelia (crista, utricle, saccule), regions (crista centre, crista periphery, utricle lateral periphery, utricle medial periphery, saccule centre, saccule external periphery, saccule internal periphery), and cell type (calyx-only HCI, HCI, HCII). Selected graphs are also shown in Fig 3. Thus, utricle data in Fig 3A-D clearly illustrate that HCI (Fig 3A, B) are more sensitive than HCII (Fig 3C, D) and that the lateral periphery (Fig 3A, C) is more resistant than the striola region (Fig 3B, D) to IDPN toxicity. The data included in Fig 3 also illustrate the large difference in sensitivity between the centre of the crista (robust effects at 3X175 in HCI, Fig 3E, and at 3X225 in HCII, Fig 3G) and the internal periphery of the saccule (first affected group 3X225 in HCI, Fig 3F, no robust effect in HCII at any dose up to 3X300, Fig 3H).

### >3.2. Relationship between HC loss and loss of vestibular reflexes

To explore the relationship between loss of the different populations of HCs and loss of the reflexes, we plotted the individual HC counts in each cell population against the corresponding tail-lift angles and air-righting times. Selected examples are shown in Fig 4 and the complete set of graphs is provided in the supplementary Figures S4-S9. These graphs suggested that the reflex measures have different strengths in their association with the cell populations, and that these appear to be non-linear. Thus, HCI in the crista centre were lost before the tail-lift angles decreased (Fig 4A) or the air-righting times increased (Fig 4D), while in contrast most HCII remained in the internal periphery of the saccule after animals suffered major alterations in both reflexes (Fig 4C and F). Meanwhile, a cell population with intermediate sensitivity to IDPN (HCII in the utricle striola) displayed an apparent correlation with the air-righting time (Fig 4E), but less apparent with the tail-lift reflex (Fig 4B).

**Figure 4.**
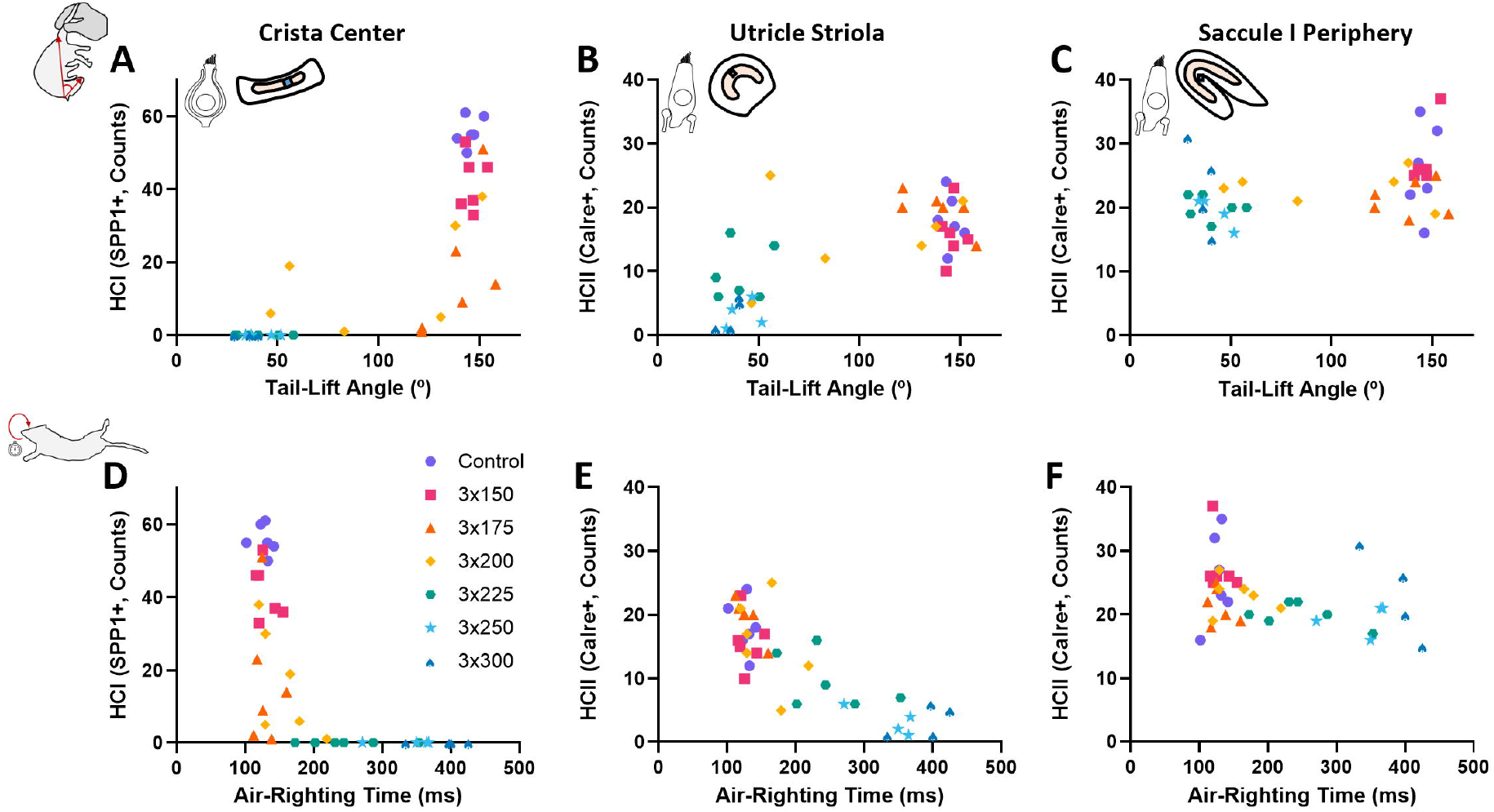
Representative examples of the relationship between HC loss and vestibular reflex alteration after exposure to 3,3’-iminodipropionitrile (IDPN) in adult female Long-Evans rats. Animals received one i.p. injection per day of IDPN (0 [Control] to 300 mg/kg/day) for 3 days. Tail-lift angles **(A-C)** and air-righting times **(D-F)** are shown paired to HCI in the crista center **(A, D)**, HCII in the utricle striola **(B, E)**, and HCII in saccule internal periphery **(C, F).** Points display data for individual animals. The legend in panel D indicating dose groups for the data point symbols applies to all panels.

To identify the underlying associations between the HC loss and the functional deficits caused by the ototoxic exposure, HC counts and behavioural measures were normalized and their relationships examined as described in the methods section. We first examined the association of cell types (HCI vs HCII) and receptor (crista, utricle, saccule) with the tail-lift reflex, using the average of the cell counts in the central and peripheral regions (Fig 5A-F). For each comparison, the extent of the Functional Cell Range (FCR, see methods) was taken as an indicator of the relevance of the association. The results suggest functional associations of the loss of the tail-lift reflex with the loss of HCI in the crista (FCR=46, Fig 5A) and the utricle (FCR=46, Fig 5B). In contrast, no clear associations were found for tail-lift deficits with HCI in the saccule (FCR=2, Fig 5C), or with HCII in the crista (FCR=2, Fig 5D), utricle (FCR=2, Fig 5E), or saccule (FCR=3, Fig 5F). We also evaluated whether the association of the tail-lift deficits with the HCI loss in the crista and utricle was preferentially linked with the central/striola regions or the peripheral regions of the different organs. As shown in Fig 5G-K, the larger FCRs were found for HCI in the periphery of the crista (FCR=74, Fig 5H) followed by the lateral periphery of the utricle (FCR=59, Fig 5K). Smaller to irrelevant associations were identified in the utricle striola (FCR=27, Fig 5I), the crista centre (FCR=10, Fig 5G) and the medial periphery of the utricle (FCR=3, Fig 5J). Minor associations were also found for the saccular regions (striola FCR=2, internal periphery FCR=4, external periphery FCR=2; data not shown).

**Figure 5.**
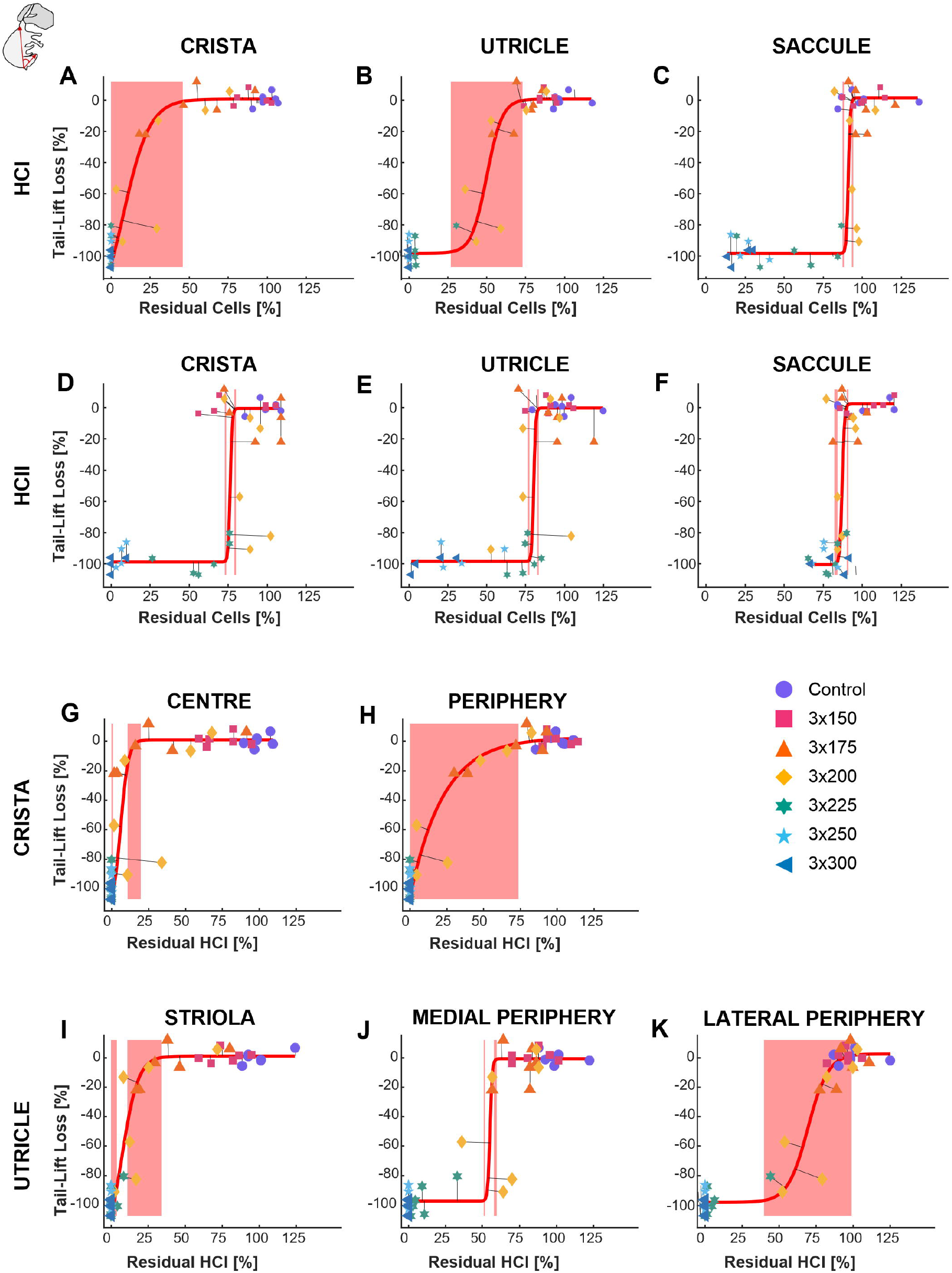
Normalized tail-lift angle as a function of the normalized number of HCs. Each data point is from one individual adult female Long-Evans rats exposed to 3,3’-iminodipropionitrile (IDPN), one i.p. injection per day at 0 [Control] to 300 mg/kg/day for 3 days as shown in the figure legend. Data in **A-F** show the average in central/striola and peripheral regions of the number of HCI **(A, B, C)** and HCII **(D, E, F)** in the crista **(A, D)**, utricle **(B, E)**, and saccule **(C, F).** In **G-K**, central/striola and peripheral data are presented separately for the crista **(G, H)** and the utricle **(I, J, K)**. Lines represent the sigmoid fits using orthogonal distance regression. Shaded areas indicate regions of the fitting curve in which the slope is greater than 10<u>°</u> and lower than 80<u>°</u>.

The association analyses with the air-righting data (Fig 6) also revealed a clearer association of the impairment of this reflex with HCI loss (Fig 6A-C) than with HCII loss (Fig 6D-F). However, in this case the association was small with the crista HCI (FCR=3, Fig 6A) and larger with the utricle HCI (FCR=52, Fig 6B) and saccule HCI (FCR =120, Fig 6C). The HCII vs air-righting reflex values were as follow: crista, FCR=8; utricle, FCR=7; and saccule, FCR=10, (Figs. 6D, 6E, and 6F, respectively). Comparing striola and peripheral regions for the association of the air-righting deficits with the HCI loss in utricle and saccule (Fig 6G-L), large associations were found for the saccule striola (FCR=110, Fig 6J) and the saccule external periphery (FCR=110, Fig 6L), followed by the utricle medial periphery (FCR=51, Fig 6H). Associations were small with the striola of the utricle (FCR=10, Fig 6G), the utricle lateral periphery (FCR=13, Fig 6I), and the internal periphery of the saccule (FCR=13, Fig 6K).

**Figure 6.**
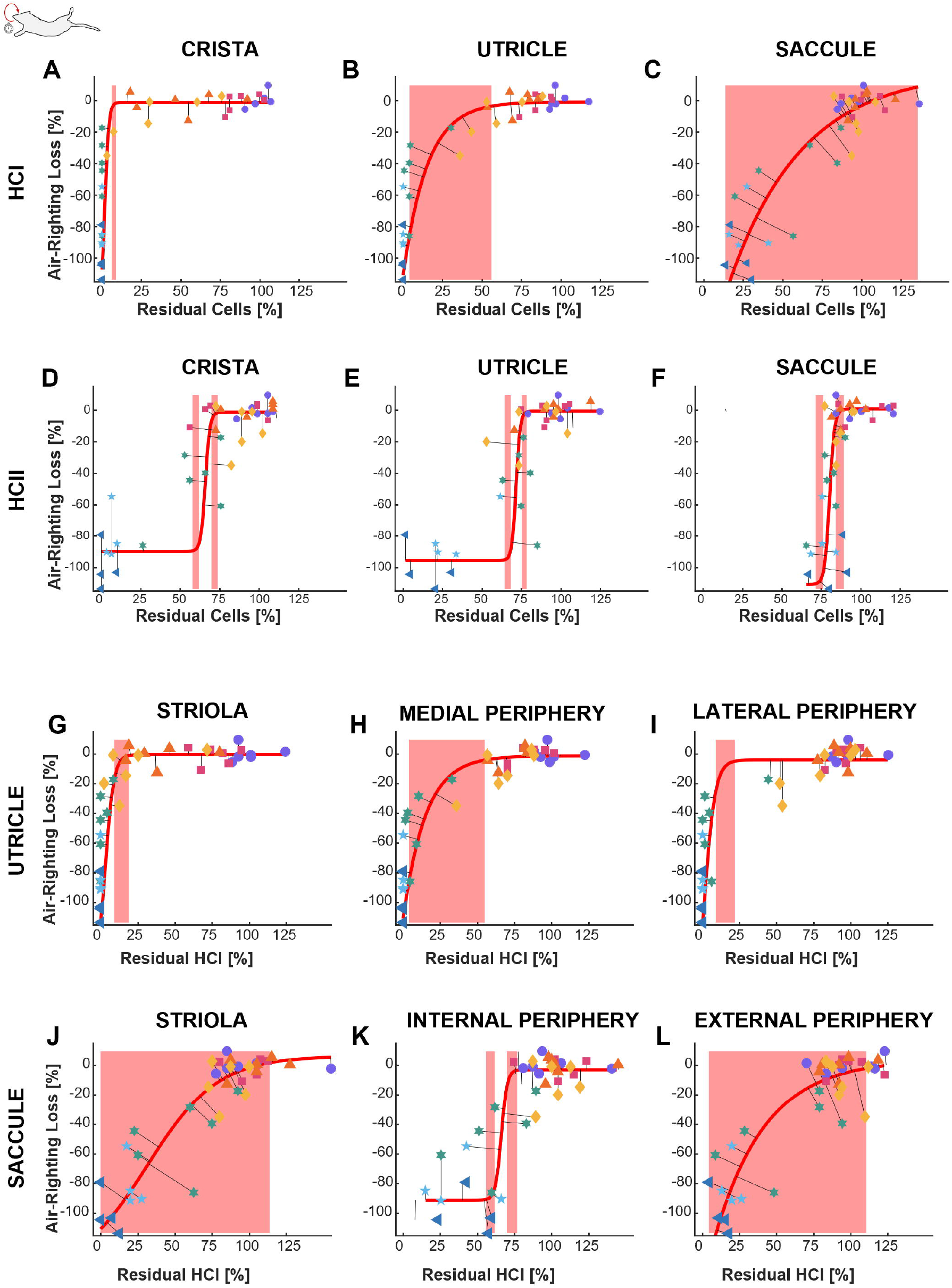
Normalized air-righting time as a function of the normalized number of HCs. Each data point is from one individual adult female Long-Evans rats exposed to 3,3’-iminodipropionitrile (IDPN), one i.p. injection per day at 0 [Control] to 300 mg/kg/day for 3 days as shown in the legend in Figure 5. Data in **A-F** show the average in central/striola and peripheral regions of the number of HCI **(A, B, C)** and HCII **(D, E, F)** in the crista **(A, D)**, utricle **(B, E)**, and saccule **(C, F).** In G-L, central/striola and peripheral data are presented separately for the utricle **(G, H, I)** and the saccule **(J, K, L)**. Lines represent the sigmoid fits using orthogonal distance regression. Shaded areas indicate regions of the fitting curve in which the slope is greater than 10<u>°</u> and lower than 80<u>°</u>.

## 4. DISCUSSION

Vestibular loss results in loss of vestibulo-motor reflexes, including the extensively studied vestibulo-ocular (VOR) reflex and the less characterized anti-gravity reflexes (tail-lift reflex, air-righting reflex). The present work continues our recent studies characterizing the impact of partial or reversible vestibular lesions caused by IDPN ototoxicity on the VOR in male and female mice (Schenberg et al., 2023; Schenberg et al., 2026) and on the anti-gravity reflexes in male rats (Martins-Lopes et al., 2019; Maroto et al., 2021a).

The toxicity of IDPN on the vestibular HCs (Llorens et al., 1993) offers a variety of animal models to characterize the functional consequences of vestibular epithelium lesions. Thus, chronic low-dose exposure causes progressive and reversible vestibular function loss in male Long-Evans rats (Llorens and Rodríguez-Farré, 1997; Sedó-Cabezón et al., 2015; Martins-Lopes et al., 2019), male 129S1/SvImJ mice (Greguske et al., 2019), and male and female C57BL/6J mice (Schenberg et al., 2023, 2026). The synaptic uncoupling observed in these chronic models (Sedó-Cabezón et al., 2015; Greguske et al., 2019; Schenberg et al., 2023) is most likely triggered by the HCs in response to the sub-lethal but persistent stress (Borrajo et al., 2025). In contrast to chronic exposure, acute or sub-acute exposure to higher doses of IDPN cause a dose-dependent irreversible loss of HCs in rodent and non-rodent species (Llorens et al., 1993; Soler-Martín et al., 2007; Wilkerson et al., 2018; Zeng et al., 2020; Maroto et al., 2021a; Schenberg et al., 2026). IDPN is known to cause also a proximal neurofilamentous axonopathy affecting large myelinated axons (Chou and Hartmann, 1964). This neuronal effect is associated with impairment of the axonal transport of neurofilaments (Griffin et al., 1978) and becomes prominent after very high acute doses (Chou and Hartmann, 1964) or after chronic exposure (Llorens and Rodríguez-Farré, 1997), but does not cause behavioural alterations that could interfere with the assessment of vestibular reflexes after exposure to the acute ototoxic doses used here (Llorens and Rodríguez-Farré, 1997; Soler-Martín et al., 2014).

In rats, Moser and Boyes (1993) described that males and females have a different dose-response relationship in their susceptibility to the behavioural syndrome elicited by IDPN, with the males being more affected. Because of this sex difference in susceptibility, conceivably related to differences on metabolic detoxification (Llorens and Crofton, 1991), most studies using IDPN in rats have included one sex only and detailed data on its vestibular toxicity are available in males only. In this work, we provide a detailed dose-response relationship of the effects of IDPN on the vestibular system of the female rat, easing the design of studies using this model in females (e.g. Chalansonnet et al., 2025). When compared to the published male data (Maroto et al., 2021a), the present data demonstrate that the reduced susceptibility of the females to its behavioural effects (Moser and Boyes, 1993) reflects a reduced susceptibility to IDPN-induced HC loss. As an example of this reduced susceptibility one can compare the data on HCI in the crista centre in females (Fig 3E) with the corresponding data in males: more than 85 % HCI loss was recorded in IDPN_3X150 males (Maroto et al., 2021a) while the same dose caused a small non-significant effect in this cell population in females. Also, all but one IDPN_3X175 and all IDPN_3X200 males had no remaining HCI (Maroto et al., 2021a), while an IDPN_3X225 dose was required to cause a similar effect in females. Besides this shift to the right in the dose-response relationship, both the female and male Long-Evans rats are suitable for detailed studies on the functional effects of the IDPN-induced vestibular HC loss.

As widely accepted but rarely measured (Lopez et al. 1997; Nakayama et al. 1996; Hirvonen et al. 2005; Maroto et al. 2021a, 2023), this study recorded a higher susceptibility to ototoxicity-induced loss of HCI with respect to HCII. In addition, there are also differences in susceptibility among organs, with the crista being slightly more sensitive to ototoxicity than the utricle and the saccule more resistant than these two organs (Fig 2; Maroto et al., 2021a). Although the vestibular system works to comprehensively encode all types and directions of head accelerations, some stimuli may trigger particular reflexes that depend exclusively (or at least almost exclusively in some circumstances) on one part of the system. This confers these reflexes high diagnostic value. For instance, the video Head Impulse Test (vHIT) is used in the clinics to independently evaluate the function of the six cristae of the patient (Curthoys et al., 2023). The differential susceptibility of the HC populations to the ototoxicity can be used to interrogate on the dependency of the reflexes on the different populations, although the analysis is complicated by the non-linear relationship relating HC loss and reflex deficits make evident by the raw data (Fig 4, Supplementary Figures S4-S9) and the existing correlations among the extent of the damage in the different cell populations (see Fig 2 in Schenberg et al., 2026). Nevertheless, this approach allowed us to obtain evidence that the tail-lift and air-righting reflexes likely depend on different vestibular HC populations (Maroto et al., 2023). In the present work, some conclusions could be drawn from the raw data graphs. For instance, with the exception of the saccule and the air-righting reflex, the calyx-only subpopulations of HCs showed scant relationship with the loss of the reflexes. However, the non-linear model analyses on the normalized data better illuminated the strength of the association of the cell types with the reflexes. Thus, the results in Figures 5 and 6 provide strong support to the hypothesis that both the tail-lift and the air-righting reflexes are triggered by HCI rather than HCII. This conclusion parallels the evidence that HCI, not HCII, have a decisive role in VOR reflexes as evaluated in mice in our previous studies (Schenberg et al., 2023; Schenberg et al., 2026). The central role of HCI in the fast vestibular reflexes is congruent with the evidence that the calyx is an adaptation supporting non-quantal synaptic transmission that is faster than standard synaptic transmission (Govindaraju et al., 2023). The analyses of our data also support the conclusions that the tail-lift reflex depends mostly on crista or utricle, not saccule, function (Fig 5A-C), and that the air-righting depends on utricle or saccule, not crista, function (Fig 6A-C). Together with the different susceptibility of the different organs to the damage, this accounts for the shift to the right observed when comparing the dose-response curves of the air-righting reflex with respect to the tail-lift reflex seen in Figure 1C,D.

The results of this study were not completely conclusive when comparing centre/striola with peripheral regions within the receptors. Nevertheless, a stronger association with the periphery of the crista and/or lateral periphery of the utricle than with the centre/striola regions of these receptors emerged for the tail-lift reflex (Fig 5G-K). For the air-righting reflex, a strong association was found for both the striola and external periphery regions of the saccule (Fig 6J-L). The possibility of differential roles for central and peripheral HCI in different behaviours is firmly supported by recent evidence in mice that centre/striola HCI are required for head stability (Ono et al., 2026) while peripheral HCI are needed for the vestibulo-ocular reflex as well as for motor performance in the rotarod and balance beam tests (Ciani Berlingeri et al., 2026). However, it is also possible that a large population of cells, including both central and peripheral areas of the receptor, are involved in triggering the reflex (Schenberg et al., 2026).

One characteristic feature of the tail-lift reflex was its abrupt transition from normal to fully deficient responses. As previously observed in male rats (Maroto et al., 2021a), few female rats showed intermediate responses, that is, tail-lift angles between 60° and 130°. There are at least two possible explanations for this observation. One possibility is that the tail-lift reflex depends on a particular small population of HCs that degenerate at a certain level of toxicity. Alternatively, the abrupt shift may result from a threshold, all or nothing, effect, in which a narrow percentage of loss in a relatively large population of HCs shifts the response from present to absent. Clearly, further research is required to explain this observation. Compared to the tail-lift reflex loss, the air-righting reflex loss showed a more graded progression.

The tail-lift and the air-righting reflexes provide non-invasive and easily-performed test of vestibulo-motor function in rats, and the present work provides critical understanding on their HC dependency. This shall facilitate their use and fulfil their translational value. For instance, the tail-lift reflex may be particularly useful in screening for ototoxicity in preclinical safety studies (Cavero and Holzgrefe, 2017), because it may reveal loss of the most sensitive HC populations, HCI in crista and utricle, and may predict functional deficits with clinical significance, such as the loss of protective reflexes during falls resulting in increased risk of head and trunk trauma. Also, they can be used to evaluate recovery of vestibular function in animal studies of regenerative therapies. From the present data one can infer that only therapies achieving significant regeneration of functional HCI in the utricle and crista will result in significant recovery of these reflexes. Current approaches mostly achieve only partial HCII regeneration (Sayyid et al., 2019; Wang et al., 2019) and limited functional recovery (Lahlou et al., 2024).

In conclusion, the present work studied the relationship between HC loss and vestibulo-spinal reflex deficits in female Long-Evans rats exposed to different doses of an ototoxic compound, IDPN, that result in graded lesions. The data strongly support the conclusion that both the tail-lift reflex and the air-righting reflex are triggered by HCI, not HCII. They also support the conclusions that the cristae and/or the utricle drive the tail-lift reflex, while the utricle and/or the saccule drive the air-righting reflex. These conclusions strengthen the use of these vestibulo-spinal reflexes in the study of vestibular dysfunction and experimental therapies in rats.

## Supporting information

Legends for Supplementary Figures

## Acknowledgments

The confocal microscopy studies were performed at the Centres Científics i Tecnològics de la Universitat de Barcelona (CCiTUB).

## Co-author contributions

A.P.: Conceptualization, Data curation, Formal analysis, Investigation, Visualization, Writing – review and editing. M.T.: Conceptualization, Formal analysis, Methodology, Visualization, Writing – review and editing.

M.B.: Conceptualization, Funding acquisition, Project administration, Writing – review and editing.

J.L.: Conceptualization, Data curation, Formal analysis, Funding acquisition, Investigation, Project administration, Supervision, Visualization, Writing – original draft, Writing – review and editing.

## Funding

This work was supported by the ERANET NEURON Program VELOSO (grants ANR-20-NEUR-0005 from Agence Nationale de la Recherche and PCI2020-120681-2 from MCIN/AEI/10.13039/501100011033 and NextGenerationEU/ PRTR) and grant PID2024-155443OB-I00 funded by MICIU/AEI/10.13039/501100011033 and European Regional Development Fund/European Union.

## Declaration of interests

The authors declare no competing interests.

## Data Availability Statement

All data supporting the results presented in the manuscript are included in the manuscript Figures

## Notes

### Competing Interest Statement

The authors have declared no competing interest.

### Summary of Updates

Authors affiliations arranged. Changed terminology to name saccular areas in text and figures. A couple of new issues included in the discussion.

