## Supplementary Figures S1-S9 for "Deficits in tail-lift and air-righting reflexes in rats after ototoxicity associate with loss of vestibular type I hair cells"

### Slide 1
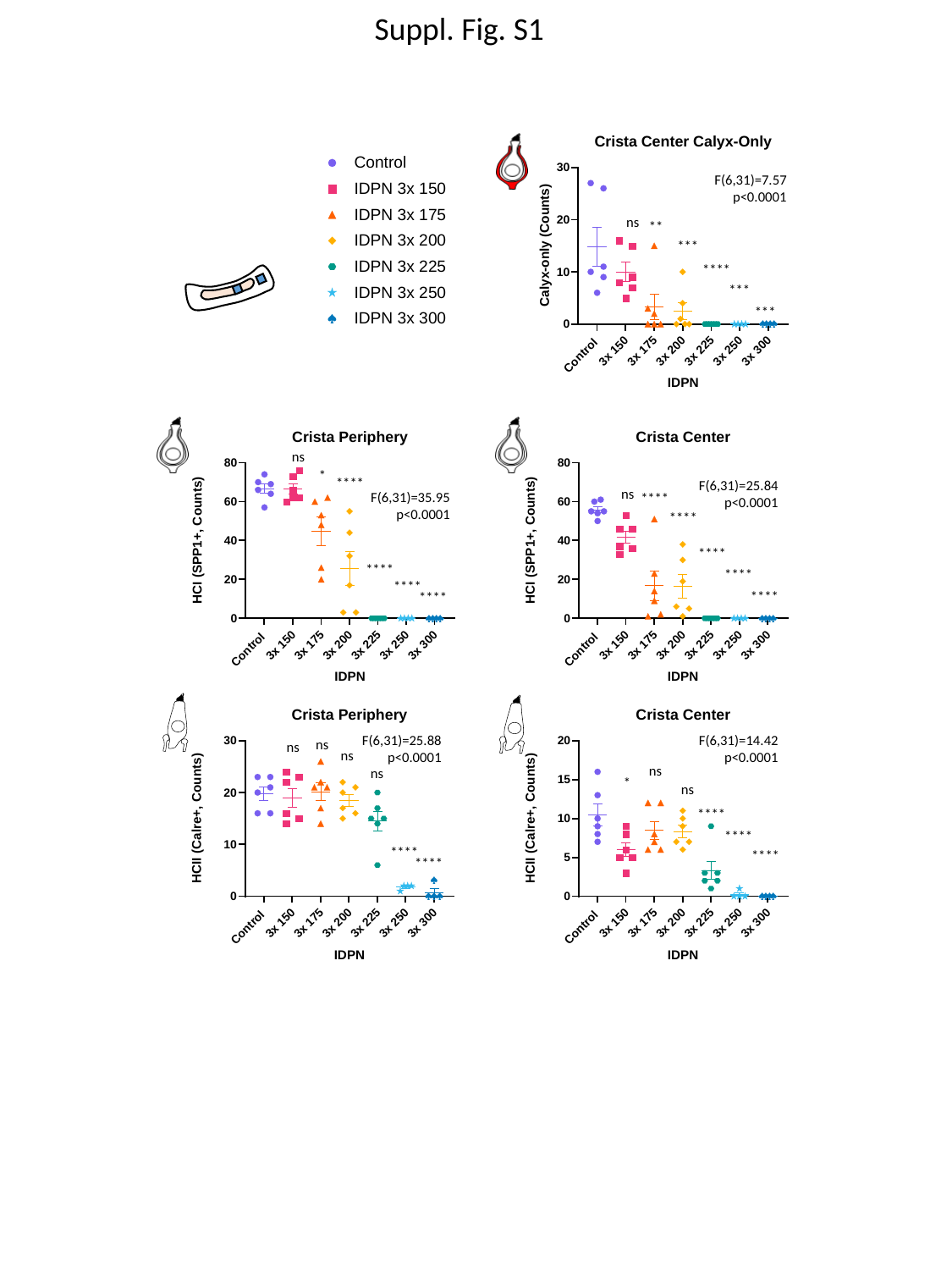

Suppl. Fig. S1
F(6,31)=7.57
p<0.0001
ns
 **
 ***
 ****
 ***
 ***
ns
*
****
F(6,31)=25.84
p<0.0001
ns
F(6,31)=35.95
p<0.0001
****
****
****
****
****
****
****
****
F(6,31)=14.42
p<0.0001
F(6,31)=25.88
p<0.0001
ns
ns
ns
ns
ns
*
ns
****
****
****
****
****

### Slide 2
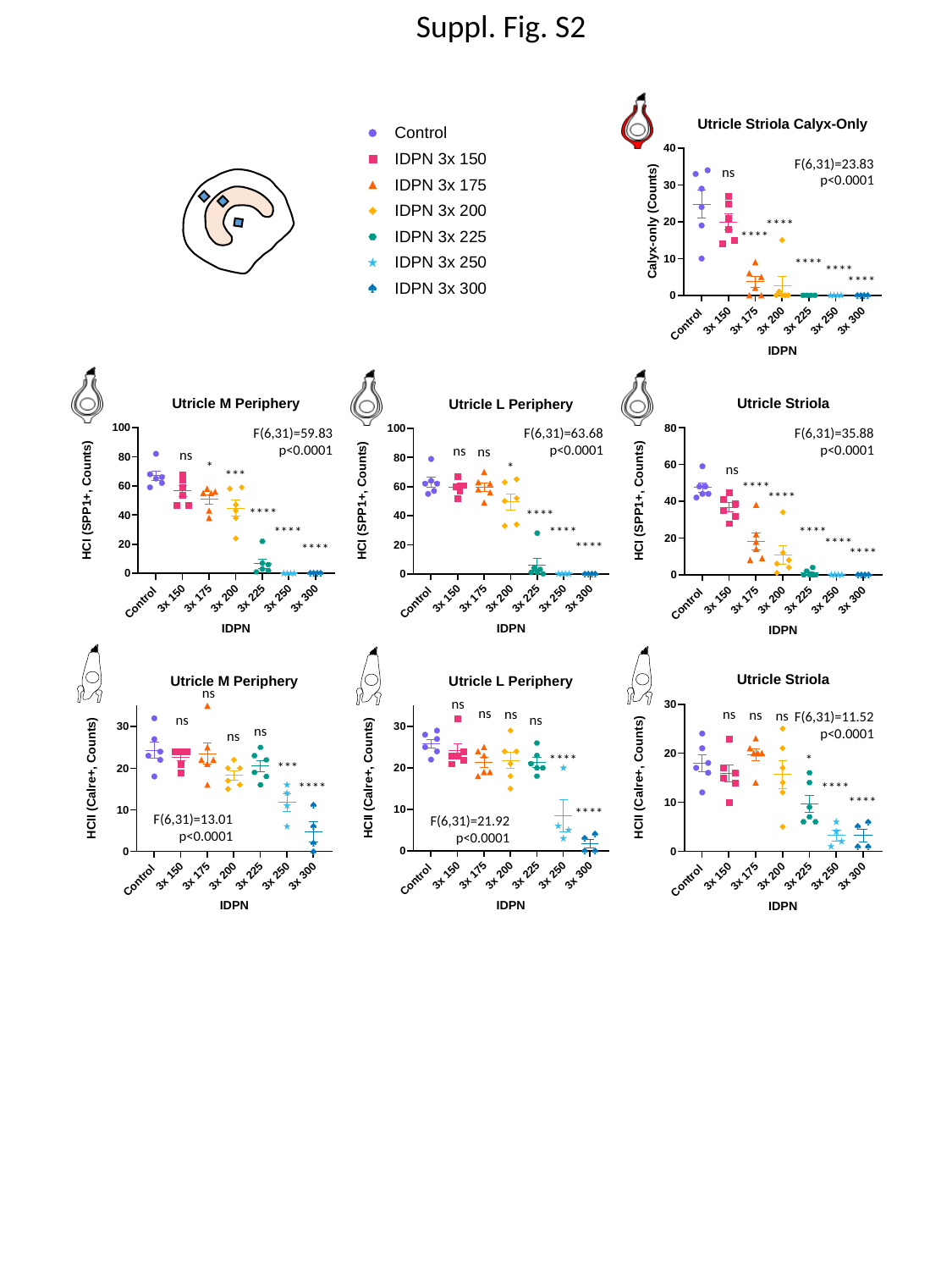

Suppl. Fig. S2
F(6,31)=23.83
p<0.0001
ns
****
****
****
****
****
F(6,31)=59.83
p<0.0001
F(6,31)=63.68
p<0.0001
F(6,31)=35.88
p<0.0001
ns
ns
ns
*
*
ns
***
****
****
****
****
****
****
****
****
****
****
****
ns
ns
ns
ns
ns
ns
ns
F(6,31)=11.52
p<0.0001
ns
ns
ns
ns
****
*
***
****
****
****
****
F(6,31)=13.01
p<0.0001
F(6,31)=21.92
p<0.0001

### Slide 3
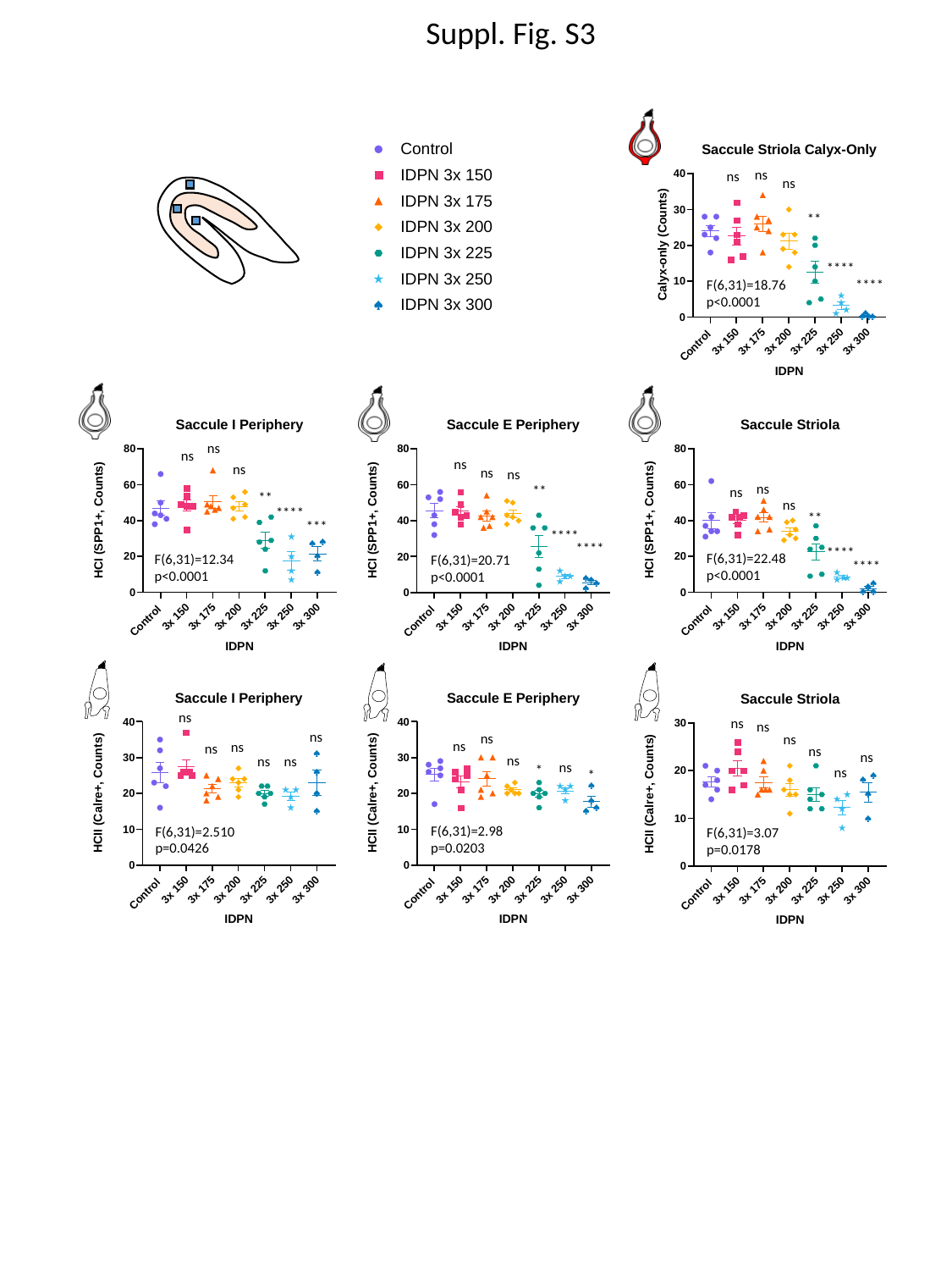

Suppl. Fig. S3
ns
ns
ns
**
****
****
F(6,31)=18.76
p<0.0001
ns
ns
ns
ns
ns
ns
ns
**
ns
**
ns
****
**
***
****
****
****
F(6,31)=22.48
p<0.0001
F(6,31)=12.34
p<0.0001
F(6,31)=20.71
p<0.0001
****
ns
ns
ns
ns
ns
ns
ns
ns
ns
ns
ns
ns
ns
ns
ns
*
ns
*
F(6,31)=2.98
p=0.0203
F(6,31)=2.510
p=0.0426
F(6,31)=3.07
p=0.0178

### Slide 4
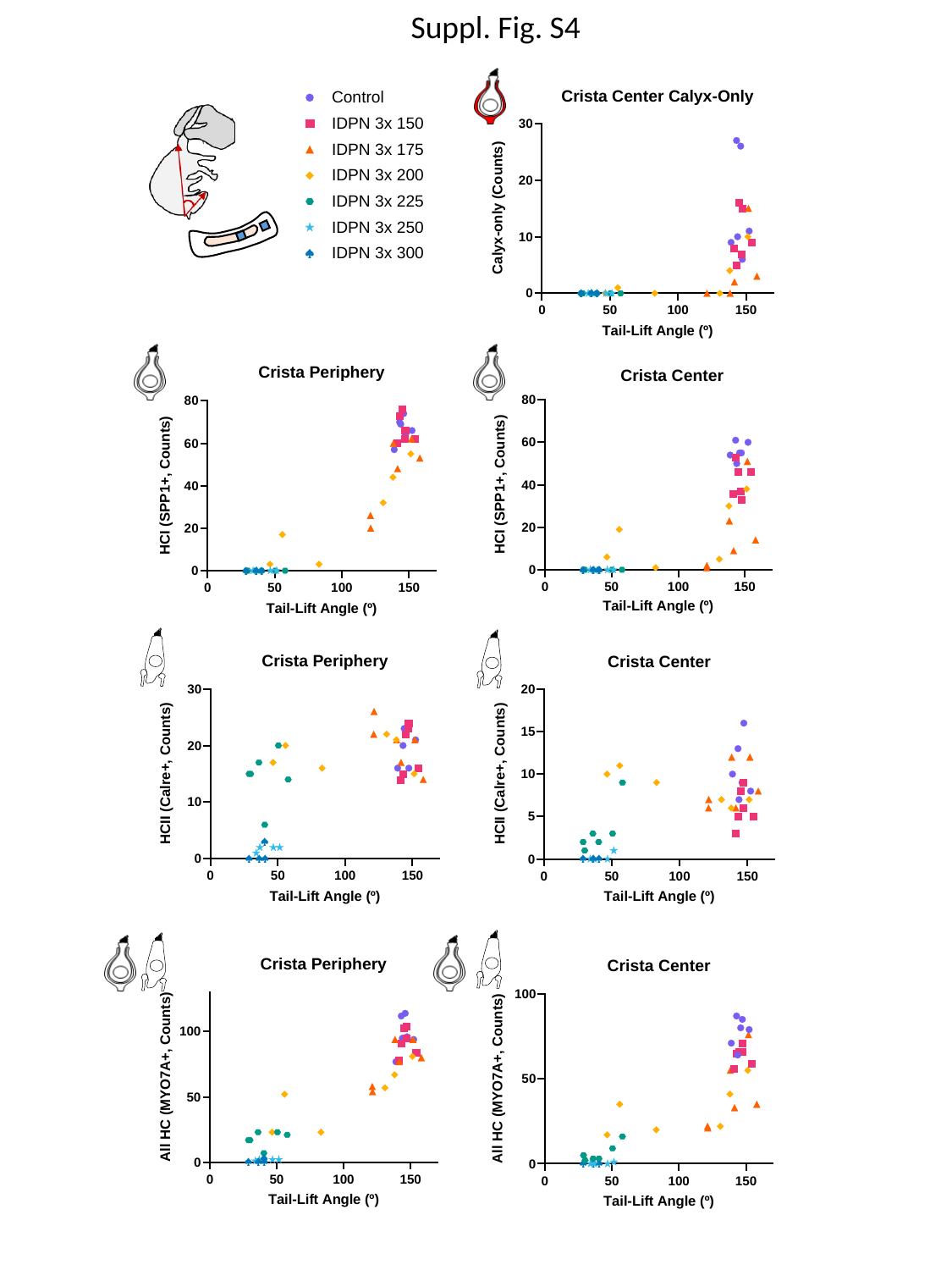

Suppl. Fig. S4

### Slide 5
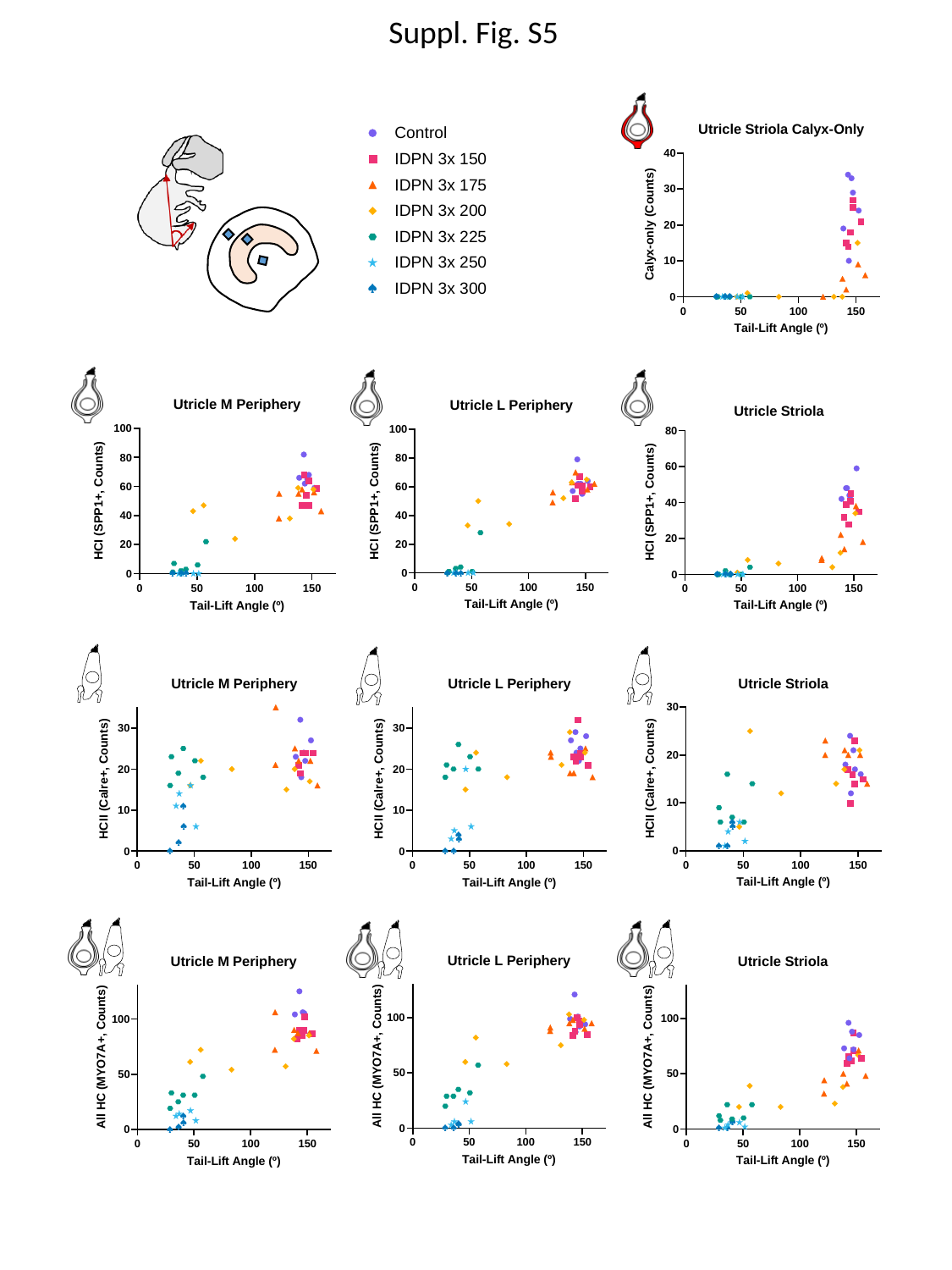

Suppl. Fig. S5

### Slide 6
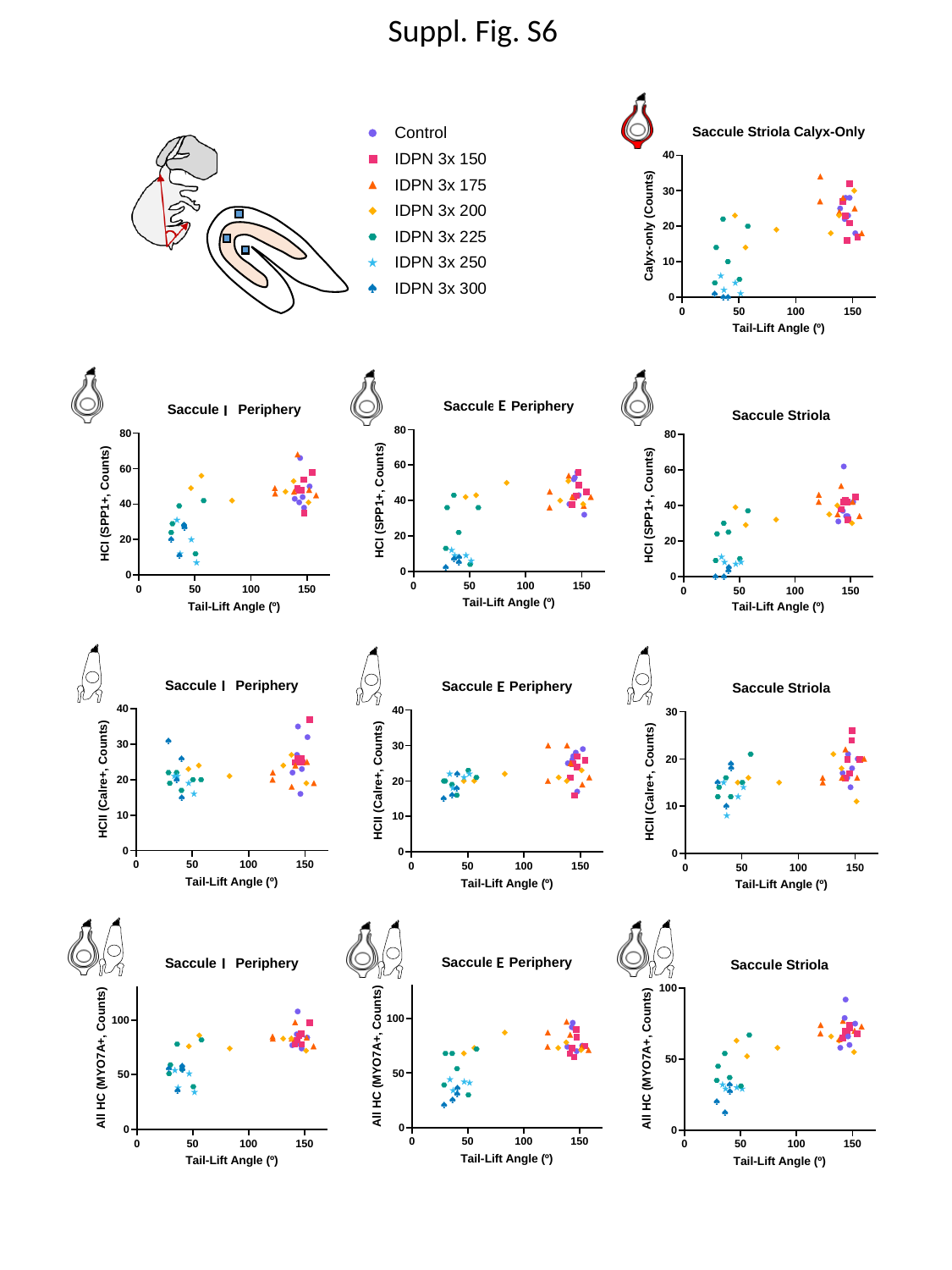

Suppl. Fig. S6
E
I
I
E
E
I

### Slide 7
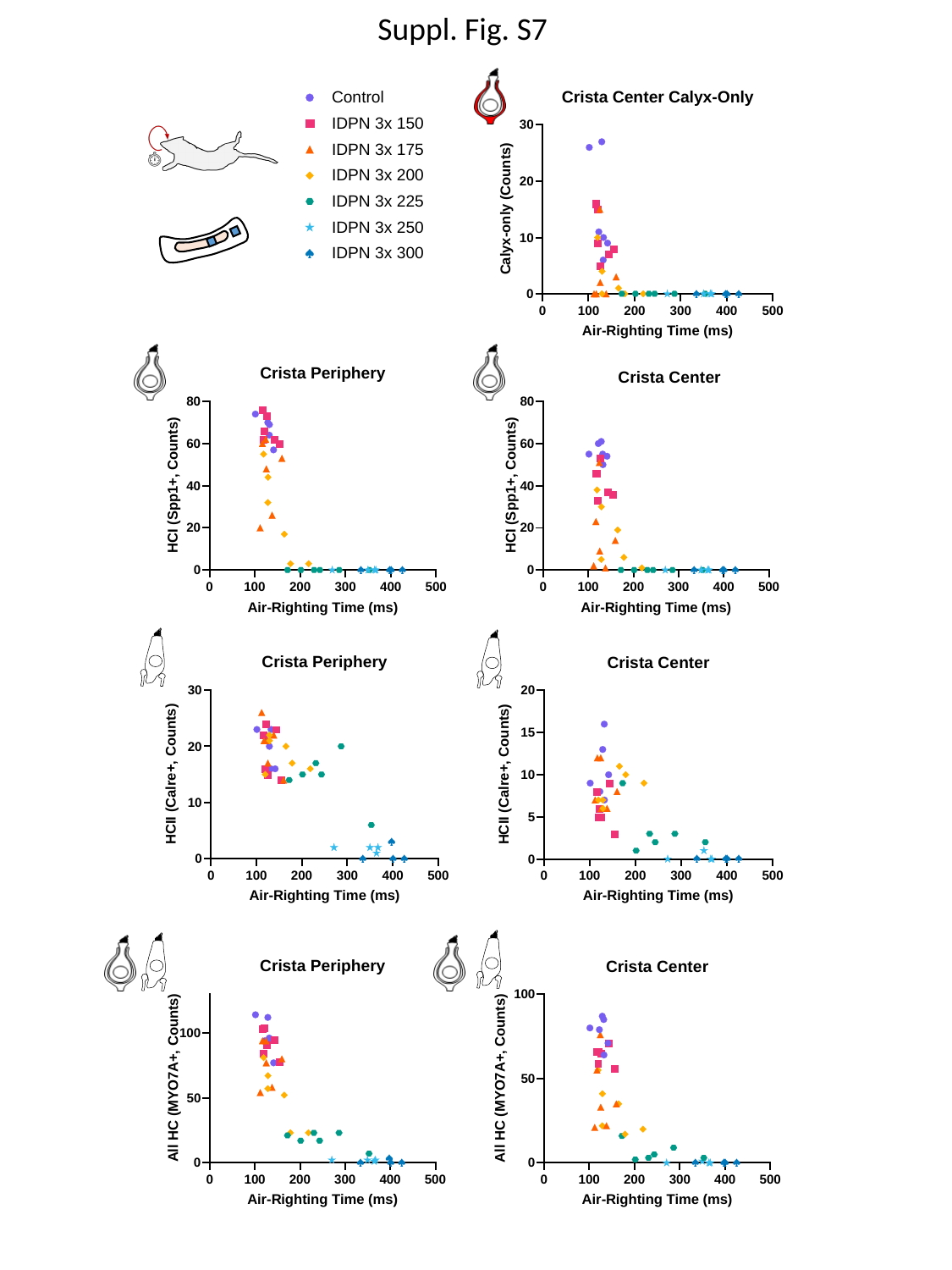

Suppl. Fig. S7

### Slide 8
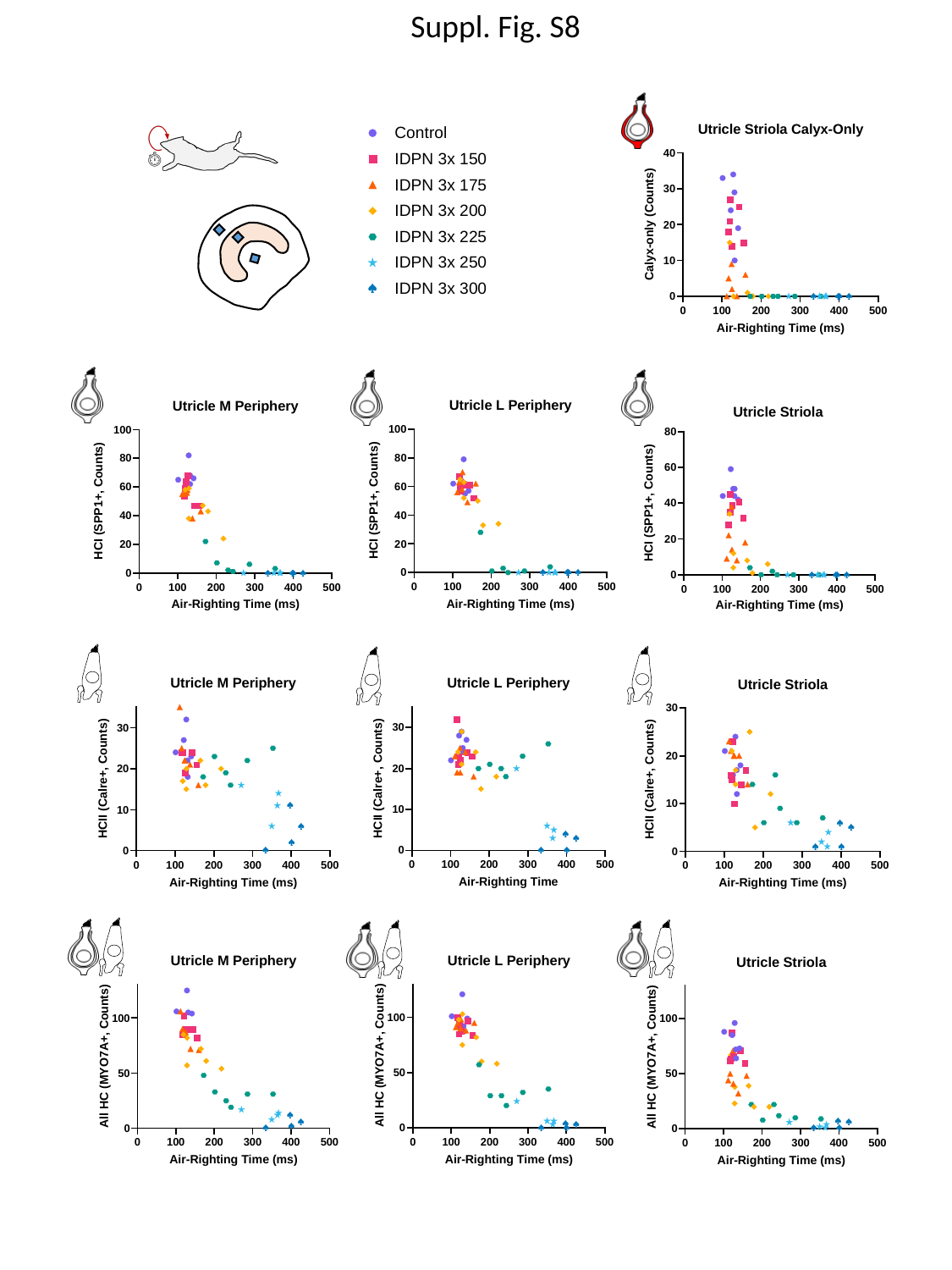

Suppl. Fig. S8

### Slide 9
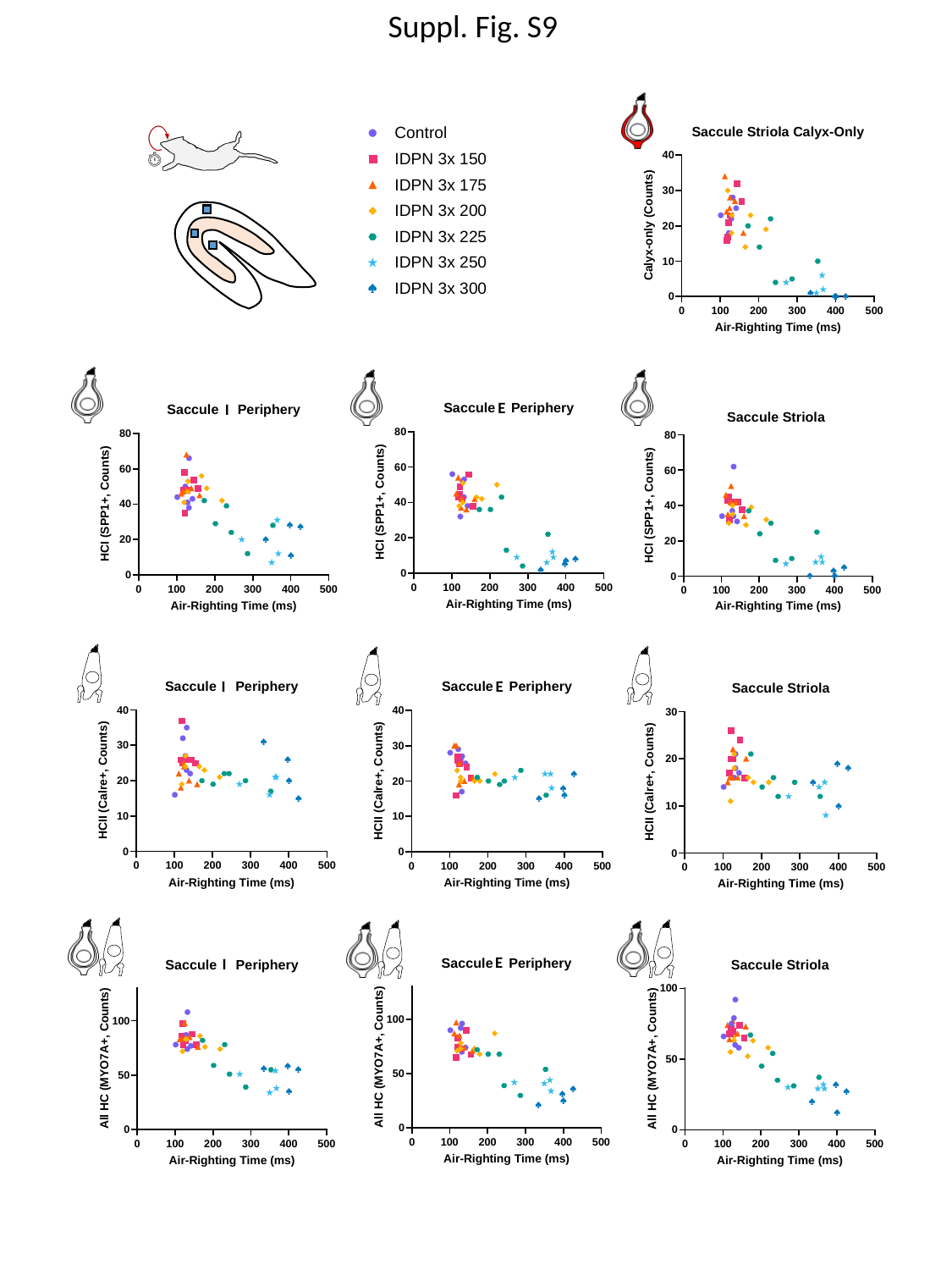

Suppl. Fig. S9
E
I
I
E
E
I
