## Supplementary material for "Deficits in tail-lift and air-righting reflexes in rats after ototoxicity associate with loss of vestibular type I hair cells": Legends for Supplementary Figures

**Palou et al. Supplementary Figure Legends**

**Figure S1**. Effects of 3,3’-iminodipropionitrile (IDPN) on hair cell (HC) counts in the vestibular crista of adult female Long-Evans rats at 26-28 days post-exposure. Counts were obtained for calyx-only-HCI, HCI, and calretinin+-HCII in the central and peripheral regions of the crista. Calyx-only-HCI are only found in the central region. Animals received one i.p. injection per day of IDPN (0 [Control] to 300 mg/kg/day) for 3 days as indicated in the abscissa. Points and lines display individual and mean ± SE values. F and p values are from one-way ANOVA analyses. Ns: p>0.05, *: p<0.05; **: p<0.01, ***: p<0.001, ****: p<0.0001, comparison to control group, Dunnett’s multiple comparisons test.

**Figure S2**. Effects of 3,3’-iminodipropionitrile (IDPN) on hair cell (HC) counts in the vestibular utricle of adult female Long-Evans rats at 26-28 days post-exposure. Counts were obtained for calyx-only-HCI, HCI, and calretinin+-HCII in the striola, lateral periphery and medial periphery regions of the utricle. Calyx-only-HCI are only found in the striola region. Animals received one i.p. injection per day of IDPN (0 [Control] to 300 mg/kg/day) for 3 days as indicated in the abscissa. Points and lines display individual and mean ± SE values. F and p values are from one-way ANOVA analyses. Ns: p>0.05, *: p<0.05; **: p<0.01, ***: p<0.001, ****: p<0.0001, comparison to control group, Dunnett’s multiple comparisons test.

**Figure S3**. Effects of 3,3’-iminodipropionitrile (IDPN) on hair cell (HC) counts in the vestibular saccule of adult female Long-Evans rats at 26-28 days post-exposure. Counts were obtained for calyx-only-HCI, HCI, and calretinin+-HCII in the striola, external periphery and internal periphery regions of the saccule. Calyx-only-HCI are only found in the striola region. Animals received one i.p. injection per day of IDPN (0 [Control] to 300 mg/kg/day) for 3 days as indicated in the abscissa. Points and lines display individual and mean ± SE values. F and p values are from one-way ANOVA analyses. Ns: p>0.05, *: p<0.05; **:p<0.01, ***:p<0.001, ****:p<0.0001, comparison to control group, Dunnett’s multiple comparisons test.

**Figure S4**. Effects of 3,3’-iminodipropionitrile (IDPN) on hair cell (HC) counts in the vestibular crista of adult female Long-Evans rats at 26-28 days post-exposure and on the tail-lift angle before tissue obtention (average of values at days 21 and 25). Animals received one i.p. injection per day of IDPN (0 [Control] to 300 mg/kg/day) for 3 days. Each point displays the relationship between the behavioural and histological data from an individual animal. Shapes and colours indicate the dose group. Calyx-only are HCIs (SPP1+) encased by a calretinin+ calyx. HCIs are SPP1+ cells. HCII are calretinin+ cells, a population comprising 85-90 % of the total HCII. All HCs are MYO7A+ cells.

**Figure S5.** Effects of 3,3’-iminodipropionitrile (IDPN) on hair cell (HC) counts in the vestibular utricle of adult female Long-Evans rats at 26-28 days post-exposure and on the tail-lift angle before tissue obtention (average of values at days 21 and 25). Animals received one i.p. injection per day of IDPN (0 [Control] to 300 mg/kg/day) for 3 days. Each point displays the relationship between the behavioural and histological data from an individual animal. Shapes and colours indicate the dose group. Calyx-only are HCIs (SPP1+) encased by a calretinin+ calyx. HCIs are SPP1+ cells. HCII are calretinin+ cells, a population comprising 85-90 % of the total HCII. All HCs are MYO7A+ cells.

**Figure S6**. Effects of 3,3’-iminodipropionitrile (IDPN) on hair cell (HC) counts in the vestibular saccule of adult female Long-Evans rats at 26-28 days post-exposure and on the tail-lift angle before tissue obtention (average of values at days 21 and 25). Animals received one i.p. injection per day of IDPN (0 [Control] to 300 mg/kg/day) for 3 days. Each point displays the relationship between the behavioural and histological data from an individual animal. Shapes and colours indicate the dose group. Calyx-only are HCIs (SPP1+) encased by a calretinin+ calyx. HCIs are SPP1+ cells. HCII are calretinin+ cells, a population comprising 85-90 % of the total HCII. All HCs are MYO7A+ cells.

**Figure S7**. Effects of 3,3’-iminodipropionitrile (IDPN) on hair cell (HC) counts in the vestibular crista of adult female Long-Evans rats at 26-28 days post-exposure and on the air-righting time before tissue obtention (average of values at days 21 and 25). Animals received one i.p. injection per day of IDPN (0 [Control] to 300 mg/kg/day) for 3 days. Each point displays the relationship between the behavioural and histological data from an individual animal. Shapes and colours indicate the dose group. Calyx-only are HCIs (SPP1+) encased by a calretinin+ calyx. HCIs are SPP1+ cells. HCII are calretinin+ cells, a population comprising 85-90 % of the total HCII. All HCs are MYO7A+ cells.

**Figure S8**. Effects of 3,3’-iminodipropionitrile (IDPN) on hair cell (HC) counts in the vestibular utricle of adult female Long-Evans rats at 26-28 days post-exposure and on the air-righting time before tissue obtention (average of values at days 21 and 25). Animals received one i.p. injection per day of IDPN (0 [Control] to 300 mg/kg/day) for 3 days. Each point displays the relationship between the behavioural and histological data from an individual animal. Shapes and colours indicate the dose group. Calyx-only are HCIs (SPP1+) encased by a calretinin+ calyx. HCIs are SPP1+ cells. HCII are calretinin+ cells, a population comprising 85-90 % of the total HCII. All HCs are MYO7A+ cells.

**Figure S9**. Effects of 3,3’-iminodipropionitrile (IDPN) on hair cell (HC) counts in the vestibular saccule of adult female Long-Evans rats at 26-28 days post-exposure and on the the air-righting time before tissue obtention (average of values at days 21 and 25). Animals received one i.p. injection per day of IDPN (0 [Control] to 300 mg/kg/day) for 3 days. Each point displays the relationship between the behavioural and histological data from an individual animal. Shapes and colours indicate the dose group. Calyx-only are HCIs (SPP1+) encased by a calretinin+ calyx. HCIs are SPP1+ cells. HCII are calretinin+ cells, a population comprising 85-90 % of the total HCII. All HCs are MYO7A+ cells.
